# A 7T interleaved fMRS and fMRI study on visual contrast dependency in the human brain

**DOI:** 10.1101/2023.07.17.548989

**Authors:** Anouk Schrantee, Chloe Najac, Chris Jungerius, Wietske van der Zwaag, Saad Jbabdi, William T Clarke, Itamar Ronen

## Abstract

**Introductions:** Functional magnetic resonance spectroscopy (fMRS) is a non-invasive technique for measuring dynamic changes in neurometabolites. While previous studies have observed concentration changes in metabolites during neural activation, the relationship between neurometabolite response and stimulus intensity and timing requires further investigation. To address this, we conducted an interleaved fMRS and functional magnetic resonance imaging (fMRI) experiment using a visual stimulus with varying contrast levels.

**Methods:** A total of 20 datasets were acquired on a 7T MRI scanner. The visual task consisted of two STIM blocks (30s/20s ON/OFF, four minutes), with 10% or 100% contrast, interleaved with a four minutes REST block. A dynamic fitting approach was used for fMRS data analysis. For metabolite level changes, the STIM conditions were modeled in two different ways: either considering the full STIM block as active condition (full-block model) or only modeling the ON blocks as active condition (sub-block model). For linewidth changes due to the BOLD effect, STIM conditions were modeled using the sub-block model.

**Results:** For both models, we observed significant increases in glutamate levels for both the 10% and 100% visual contrasts, but no significant difference between the contrasts. Decreases in aspartate, and glucose, and increases in total N-acetylaspartate and total creatine were also detected, although less consistently across both 10% and 100% visual contrasts. BOLD-driven linewidth decreases and fMRI-derived BOLD increases within the MRS voxel were observed at both 10% and 100% contrasts, with larger changes at 100% compared to 10% in the fMRI-derived BOLD only. We observed a non-linear relation between visual contrast, the BOLD response, and the glutamate response.

**Conclusion:** Our study highlights the potential of fMRS as a complementary technique to BOLD fMRI for investigating the complex interplay between visual contrast, neural activity, and neurometabolism. Future studies should further explore the temporal response profiles of different neurometabolites and refine the statistical models used for fMRS analysis.

## Introduction

Functional magnetic resonance spectroscopy (fMRS) is a powerful non-invasive technique that measures dynamic changes in neurometabolites. Unlike fMRI, which relies on hemodynamic coupling, fMRS provides a more proximal measure of neural activity. fMRS allows time-resolved measurements of metabolite concentration while the subject performs a mental task or responds to a stimulus. This makes it possible to examine the temporal behavior of metabolite levels involved in neuroenergetics and neurotransmission in human subjects.

Previous studies have demonstrated that stimulus-induced increases in neural activity result in concentration changes in certain metabolites (for reviews, see Koush et al., Mullins et al., Stanley et al.). For example, visual stimulation resulted in increases in lactate (Lac) and glutamate (Glu), but also in decreases in glucose (Glc) and aspartate (Asp) in the primary visual cortex (V1) (e.g. Prichard et al., 1991; Mangia et al., 2007; Schaller et al., 2013). Moreover, motor tasks, painful stimuli and even higher-order cognitive paradigms have also been found to elicit changes in cortical Glu and γ-aminobutyric acid (GABA) levels (e.g. Floyer-Lea et al., 2006; Mullins et al., 2005; Volovyk & Tal, 2021; Koush et al., 2021). In the initial development of fMRS task paradigms, prolonged sensory stimuli lasting several minutes were typically employed. This was partially related to the need for multiple spectral averages to ensure sufficient signal-to-noise ratio (SNR) for the stimulation and rest blocks. Moreover, changes in neurometabolite levels were hypothesized to be the result of increased oxidative metabolism to a new steady state level (Mangia et al., 2007). This supported the use of sustained stimulation as considerable time was needed for these changes to become discernible. Yet, with the development of higher magnetic fields, increased SNR allowed for better spectral resolution and improved temporal resolution (Mlynárik et al., 2008; Tkác et al., 2009; Ladd et al., 2018). As such, more recent studies have also adopted shorter block paradigms (e.g. Ip et al. 2017; Ip et al. 2019), more in line with fMRI stimulus paradigms and event-related experiments (e.g. Apsvalka et al., 2014; Stanley et al., 2017; Koolschijn et al., 2023). This approach has the advantage of reducing the habituation of the neural response, as well as allowing for a more detailed investigation of the temporal dynamics of fMRS.

Despite these developments, the exact temporal response profile for fMRS remains to be determined, highlighting the importance of further research in this area. For example, different neurometabolites may exhibit distinct temporal behavior, and different stimuli may elicit a unique temporal response. Additionally, the relation between neurometabolite response and stimulus intensity has yet to be fully characterized. It is hypothesized that, similarly to fMRI (Boynton et al., 1996; Horner & Andrews, 2009), the fMRS response will be linearly correlated with neural activity over a certain range. However, Ip et al. (2019) showed a linear relationship with image contrast for Glu and for the BOLD signal changes, using a 60-second visual stimulation paradigm. Characterizing the temporal and intensity-dependent response profiles of fMRS is crucial for a better understanding of the neural correlates of the changes observed in neurometabolite levels in fMRS.

To characterize the contrast dependency and the role of experiment timings in fMRS, we conducted an interleaved fMRS and fMRI experiment using a full-field flashing checkerboard visual stimulus. The experiment consisted of two stimulation blocks (STIM), with visual contrast levels of 10% and 100% respectively, that were alternated with REST blocks. To limit habituation to the stimulus, each long STIM block was subdivided into shorter ON-OFF periods (30s on, 20s off). This approach enabled us to analyze the temporal response function of neurometabolites by examining two time scales: the longer full STIM block (ON and OFF period included) and the shorter ON STIM blocks only. We employed a recently developed dynamic fitting approach for the analysis of our fMRS data (Clarke et al., 2023) and evaluated the contrast dependence for both fMRS and fMRI data.

## Methods

### Subjects

Subjects were scanned in two sites: at the Leiden University Medical Center (LUMC) and the Spinoza Centre for Neuroimaging. A total of 20 datasets were obtained from 19 healthy subjects (4 men and 15 women aged 21 to 43; one subject was scanned at both locations). The study adhered to the guidelines from the Institutional Review Board of the AMC and the LUMC (the Netherlands). Written informed consent was obtained from all subjects prior to the study. All were healthy normal participants who had normal or corrected-to-normal vision and had no 7T MRI contraindications. No additional screening for neurological or psychiatric illness was performed.

### MR acquisition

Data were acquired on two similar 7T MR systems (Achieva, Philips, Best, the Netherlands). A head coil consisting of a quadrature birdcage transmit and 32-channel phased-array receive coils (Nova Medical Inc., Wilmington, MA, USA) was used for all measurements. A T1-weighted (T1w) structural scan was acquired for each participant (TR/TE = 5/2 ms; FOV(AP,FH,RL) = 246×246×180 mm^3^; voxel size = 0.85×0.85×1 mm^3^; flip angle = 7°). fMRS and fMRI were interleaved with a combined dynamic scan time of 5 s (Henningsson et al., 2015). fMRS data were acquired using a sLASER sequence with FOCI refocusing pulses (Arteaga de Castro et al., 2013) and VAPOR water suppression (TR/TE = 3600/36 ms; bandwidth = 3kHz; 1024 data-points; volume-of-interest (VOI) = 14×31×14 mm^3^). Two unsuppressed water spectra were obtained for eddy current correction. The VOI was placed in the primary visual cortex (V1), based on a short checkerboard localizer fMRI sequence and identification of the calcarine sulcus on the T1w image (Figure 1B). Two outer volume suppression bands were carefully placed to suppress signal from outside the VOI, particularly the >extracerebral fat signal. fMRI data were acquired using a 3D-EPI sequence (TR/TE/FA = 31/20ms/10°; FOV(AP,FH,RL) = 240×136×240 mm^3^; voxel size = 1.875×1.875×2 mm^3^). We used dynamically alternating linear shims and a shared set of static second order terms to optimize shim settings for both the fMRS and fMRI acquisition (Boer et al., 2020). An in-house developed dielectric pad or metasurface was placed directly below the occipital cortex to maximize the transmit magnetic field (B_1_^+^) homogeneity and efficiency in the occipital regions as previously described (Webb et al. 2022).

**Figure 1.**
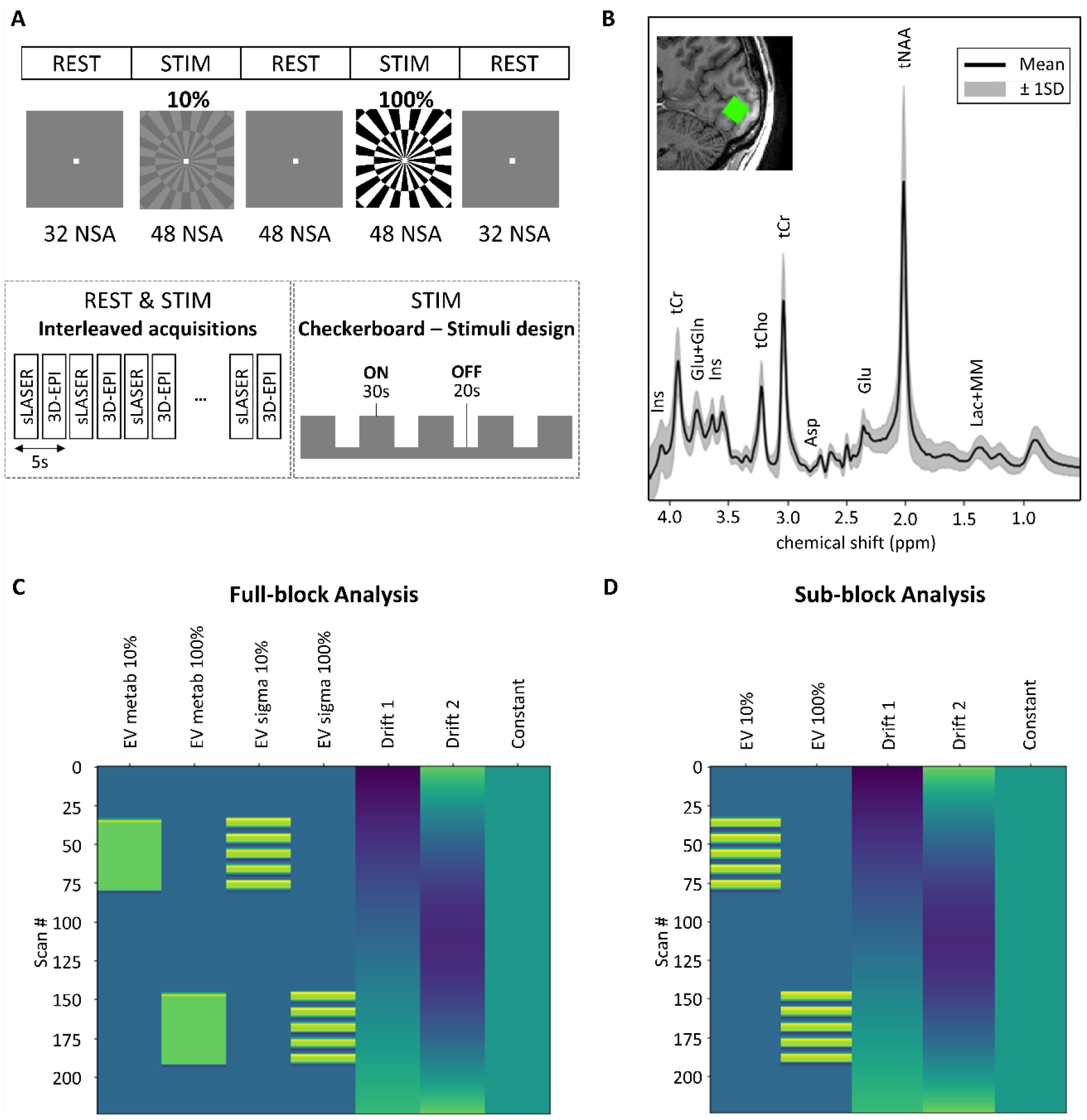
Overview of the experiment. A) schematic representation of the experiment. NSA = number of signal acquisitions per block. B) Mean spectrum across subjects averaged across transients (black line) with 1 standard deviation (gray shading). The insert shows an example voxel placement overlaid on the subject’s T1w image. C) Design matrix for the full-block analysis D) Design matrix for the sub-block analysis.

### Experimental paradigm

The task consisted of two 4-minute STIM blocks, each with different black/white contrast: 10% and 100%. The paradigm started and ended with a 2.5 min REST block and the STIM blocks were separated by a 5 min REST block. Each STIM block was subdivided into sub-blocks of 30 s ON and 20 s OFF. The total task duration was 18.5 minutes with a total of 224 individual acquisitions for both fMRS and fMRS. The stimulus consisted of a full-field radial checkerboard flickering at 8 Hz at either 10 or 100% visual contrast. A gray static image of average luminosity equal to the checkerboard with a central white fixation dot was presented during rest periods. Stimuli were generated using PsychoPy v3 (Peirce et al., 2019), viewed through a mirror mounted on the RF head coil, and displayed on a 32” BOLD screen from Cambridge Research Systems (Amsterdam) or back-projected onto a screen (Leiden). Subjects were instructed to focus on the fixation point during the entire experiment. For a schematic representation of the stimulus paradigm, see Figure 1A.

### fMRS processing and analysis

Processing steps were performed with in-house Matlab scripts (R2019b, The MathWorks, Inc. USA). Raw data were phase- and amplitude corrected before coil combination. Subsequently, data were corrected for eddy currents, and spectral registration was used to correct for frequency and phase drifts across individual acquisitions (Near et al., 2015). A basis set was simulated based on our MRS sequence in FSL-MRS (Clarke et al., 2021). The basis set was simulated using full pulse descriptions and 60 spatially resolved points in each dimension. The data were exported from Matlab and converted to NIfTI-MRS format (Clarke et al. 2022). A second alignment step was performed using FSL-MRS’s dynamic alignment tool. Briefly, this performs a full linear combination modeling fit of each time point, using the above basis set, with all parameters except global phase and frequency shift fixed across time. Subsequently the fitted phase and frequency shifts are removed from the data.

#### For quality control

For each subject, all individual transients were averaged and the average spectrum was fitted using FSL-MRS (version 2.1.6) to determine spectral quality. Briefly, basis spectra are fitted to the complex-valued spectrum in the frequency domain, using our simulated basis set, that included 19 metabolites and a measured macromolecular baseline. The basis spectra are shifted and broadened with a Voigt lineshape model with parameters fitted to the data and grouped into 2 metabolite groups (first group: metabolites, second group: macromolecules). A complex second order polynomial baseline is also concurrently fitted. Model fitting was performed using the truncated Newton algorithm as implemented in *scipy* (Virtanen et al. 2020). Metabolites were excluded from further analysis if the CRLB exceeded 20% in more than 50% of the subjects based on the average spectra. Additionally, we report the linewidth (i.e.full-width-half-maximum (FWHM)) of N-acetylaspartate (NAA) and the Cramer-Rao Lower Bounds (CRLB) of Glu as metrics for spectral quality.

#### For the dynamic analysis

All fMRS data were analyzed using a general linear model (GLM). Design matrices (created using *nilearn* (Abraham et al., 2014)) were created, with both the 10% and 100% as separate explanatory variables (EV) in the design matrix, as well as a linear drift, a quadratic drift and a baseline EV (Figure 1C,D). The STIM conditions were modeled in two different ways: either considering the full STIM block as active condition (*full-block model*, including both ON and OFF parts of STIM blocks) or only modeling the sub-blocks within the STIM block as active condition *(sub-block model*, only ON parts of STIM blocks) (Figure 1C,D). The dynamic spectral fitting model of FSL-MRS was used to assess the time-dependence of the metabolite concentration on the design matrix, using our simulated FSL-MRS basis set. The phase, shift and baseline parameters were always assumed to be fixed over time, whereas the concentrations were constrained by the model (design matrix) (Supplementary Figure 1). Previous studies have demonstrated that stimulus-induced increases in the BOLD effect can induce line-narrowing in MR spectra as a result of changes in T2* (Zhu & Chen, 2001; Mangia et al., 2006). To take these potential effects into account, we evaluated two models: a fixed linewidth model and a variable linewidth model, which additionally applies the same design matrix to the Gaussian line-broadening parameter (sigma); the Lorentzian parameter (gamma) was fixed across time. Please note that for both the full-block and sub-block models the line-broadening parameter is modeled according to the sub-block model, as this best reflects the expected hemodynamic response function (Figure 1C,D). Spectral fitting parameters are as described above in ‘for quality control’. The fMRS acquisition and analysis parameters are reported in the MRSinMRS checklist (Lin et al., 2021; Supplementary Table 1). In order to display the individual data points of the fits, we extracted the results of the initial (single transient) fits of Glu for each subject. These results were then averaged across all subjects and smoothed using a moving average with a bin size of 32.

**Table 1.** Statistical results for the full-block analysis.

|  | 10% |  | 100% |  | mean activation |  | 100 vs 10% |  |
| --- | --- | --- | --- | --- | --- | --- | --- | --- |
|  | *z* | *p* | *z* | *p* | *z* | *p* | *z* | *p* |
| **Metabolite level** |  |  |  |  |  |  |  |  |
| *Asc* | 0.98 | 0.16 | 0.32 | 0.37 | 0.60 | 0.27 | -0.46 | 0.32 |
| *Asp* | -1.75 | 0.04 | **-2.03** | **0.02** | **-2.27** | **0.01** | -0.29 | 0.39 |
| *GSH* | 0.03 | 0.49 | 1.14 | 0.13 | 0.80 | 0.21 | 0.84 | 0.20 |
| *Glu* | **2.23** | **0.01** | **3.28** | **<0.001** | **3.15** | **<0.001** | 1.27 | 0.10 |
| *Ins* | -0.74 | 0.22 | 1.75 | 0.04 | 0.70 | 0.24 | 1.64 | 0.0496 |
| *PE* | -1.12 | 0.13 | 0.74 | 0.23 | -0.66 | 0.25 | 1.27 | 0.10 |
| *Scyllo* | 0.28 | 0.39 | 0.08 | 0.47 | 0.23 | 0.41 | -0.16 | 0.44 |
| *Glc+* | -1.70 | 0.04 | **-2.07** | **0.02** | **-2.49** | **0.006** | -0.24 | 0.40 |
| *tCh* | 1.11 | 0.13 | -1.27 | 0.10 | -0.12 | 0.45 | -1.67 | 0.048 |
| *tCr* | -0.55 | 0.29 | 1.91 | 0.03 | 1.34 | 0.09 | 1.62 | 0.05 |
| *tNAA* | **-2.66** | **0.003** | 0.44 | 0.67 | -2.15 | 0.02 | -1.65 | 0.049 |
| **Linewidth** |  |  |  |  |  |  |  |  |
| *sigma* | **-3.27** | **<0.001** | **-4.62** | **<0.001** | **-4.35** | **<0.001** | -1.30 | 0.10 |

Positive and negative *z*-values indicate a neurometabolite increase and decrease, respectively, in case of the 10%, 100% and mean activation. For the contrast between 10% vs 100% a positive *z*-value indicates 100% > 10%, whereas a negative *z*-value indicates 10% > 100% (in absolute terms). The significant effects corrected for multiple comparison correction ( $p < 0.025$ ) are displayed in bold. Mean first level model fit across subjects: AIC=322433.5

### fMRI processing and analysis

FMRI data processing was carried out using FEAT Version 6.00, part of FSL (FMRIB’s Software Library, www.fmrib.ox.ac.uk/fsl). Preprocessing included removal of the first two time points, motion correction, high-pass temporal filtering (0.002 Hz) and spatial smoothing using a Gaussian kernel of 5 mm FWHM. fMRI data were co-registered to the T1w scan using boundary-based registration, and then subsequently registered to MNI space. Similar to the fMRS analyses, a design matrix (‘the sub-block model’) was created with both the 10% and 100% as separate EVs (but without the drift EVs), and fed into first-level analyses in FEAT.

### Statistical analysis fMRS and fMRI

Second level GLMs (across subjects, random effect) were used to calculate group level statistics using FSL’s *flameo* (Woolrich et al., 2004) for both the fMRS and fMRI data.

For the fMRS data, we evaluated the following statistical contrasts: a) 10% vs rest b) 100% vs rest c) mean activation vs rest (i.e. average over 10% and 100% vs rest) d) 100% vs 10%. We calculated these contrasts for all metabolites that could be fitted with sufficient certainty. For some metabolites only the combined neurometabolite level was considered, due to the high correlation between components: total NAA (*tNAA*, NAA+NAAG), total creatine (*tCr*, creatine+phosphocreatine) and total choline (*tCho*, phosphocholine+glycerophosphocholine), glucose + taurine (here referred to as *Glc+*). We ran these analyses both for the full-block model and for the sub-block model, and each using the fixed linewidth model and variable linewidth model.

For the fMRI data, we evaluated the following statistical contrasts: a) 10% vs rest b) 100% vs rest c) 100% vs 10% for the change in BOLD signal (based on the sub-block design). Additionally, we extracted the BOLD time courses and BOLD contrast estimate parameters within the MRS VOI location using *fslmeants* and *featquery* respectively.

Finally, we computed Pearson’s correlation coefficients between the interindividual parameter estimates of the 10% contrast, the 100% contrast and the 100%>10% contrast for the Glu (fMRS), sigma (fMRS) and the BOLD (fMRI) response to assess the association between these modalities. As we compared two models, i.e. full-block and sub-block, we corrected the p-value for multiple comparisons using the Bonferroni correction, resulting in a corrected ɑ of 0.025.

## Results

### Data quality of average spectra

Changes in concentrations of alanine (Ala), gamma-aminobutyric acid (GABA), glutamine (Gln), and lactate (Lac) were not statistically evaluated as their CRLB exceeded 20% in more than 50% of the subjects. A full list of the reported metabolites can be found in Table 1 and 2. The mean FWHM of NAA was 14.11 (SD=1.90), with a range of 12.28 to 19.04. Additionally, the mean CRLB of Glu in the average spectra was 4.20 % (SD=1.04 %), ranging from 3.07 % to 7.46 %.

**Table 2.** Statistical results for the sub-block analysis.

|  | 10% |  | 100% |  | mean activation |  | 100 vs 10% |  |
| --- | --- | --- | --- | --- | --- | --- | --- | --- |
|  | *z* | *p* | *z* | *p* | *z* | *p* | *z* | *p* |
| **Metabolite level** |  |  |  |  |  |  |  |  |
| *Asc* | 1.11 | 0.13 | 0.25 | 0.40 | 0.71 | 0.24 | -0.57 | 0.29 |
| *Asp* | -1.26 | 0.10 | -1.21 | 0.11 | -1.61 | 0.054 | 0.08 | 0.47 |
| *GSH* | 0.57 | 0.28 | 1.09 | 0.14 | 1.13 | 0.13 | 0.41 | 0.34 |
| *Glu* | **1.99** | **0.02** | **2.94** | **0.002** | **2.87** | **0.002** | 0.24 | 0.40 |
| *Ins* | -0.04 | 0.48 | 1.85 | 0.03 | 1.30 | 0.10 | 1.10 | 0.14 |
| *PE* | -1.00 | 0.16 | 0.96 | 0.17 | -0.33 | 0.37 | 1.22 | 0.11 |
| *Scyllo* | -0.68 | 0.25 | 0.20 | 0.42 | -0.35 | 0.36 | 0.61 | 0.27 |
| *Glc+* | -1.45 | 0.07 | -1.10 | 0.15 | -1.75 | 0.04 | 0.45 | 0.32 |
| *tCh* | 1.85 | 0.03 | 0.01 | 0.50 | 1.61 | 0.054 | -1.69 | 0.046 |
| *tCr* | 0.30 | 0.38 | **2.26** | **0.01** | **2.25** | **0.01** | 1.36 | 0.09 |
| *tNAA* | -1.18 | 0.12 | 0.82 | 0.21 | -0.01 | 0.50 | 1.15 | 0.12 |
| **Linewidth** |  |  |  |  |  |  |  |  |
| *sigma* | **-2.84** | **0.001** | **-3.24** | **<0.001** | **-3.89** | **<0.001** | -1.18 | 0.12 |

Positive and negative *z*-values indicate a neurometabolite increase and decrease, respectively, in case of the 10%, 100% and mean activation. For the contrast between 10% vs 100% a positive *z*-value indicates 100% > 10%, whereas a negative *z*-value indicates 10% > 100% (in absolute terms). The significant effects corrected for multiple comparison correction ( $p < 0.025$ ) are displayed in bold. Mean first level model fit across subjects: AIC=322449.1

### fMRS

#### Full-block model

Using the variable linewidth model, we found significant line narrowing for both the 10% and 100% contrasts (10%: z=-3.27, p<0.001; 100%: z=-4.62, p<0.001), but no difference between the contrasts (z=-1.30, p=0.10). We here report the results from the variable linewidth model, whereas the results of the fixed linewidth model can be found in Supplementary Table 2. We found a significant effect of visual stimulation on Glu levels for both the 10% and 100% contrast (10%: z=2.23, p=0.01; 100%: z=3.28, p<0.001), as well as across both contrasts (z=3.15, p<0.001). We found no significant difference in Glu levels between the 10% and 100% contrasts (z=1.27, p=0.10). Decreased neurometabolite concentrations during visual stimulation were observed for aspartate (Asp) and Glc+ during the 100% contrast (Asp: z=-2.03, p=0.02; Glc+: z=-2.07, p=0.02) and across STIM blocks (mean activation Asp: z=-2.27, p=0.01; Glc+: z=-2.49, p=0.006), and tNAA during the 10% contrast (z=-2.66, p=0.003). These results are displayed in Figure 2 and Supplementary Figure 2. All statistical results can be found in Table 1.

**Figure 2.**
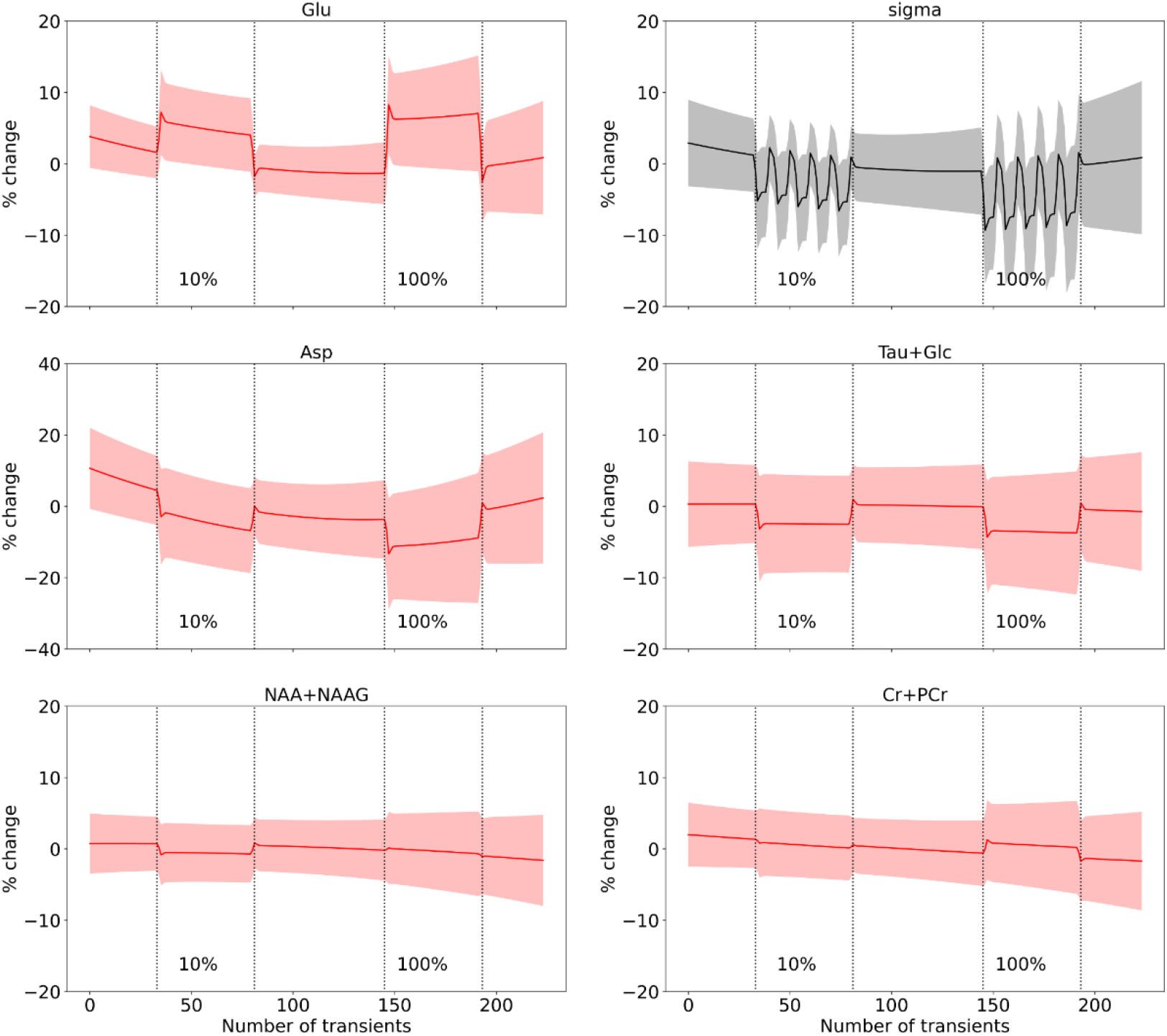
fMRS full block results. The figure displays significant results from the second-level fMRS analysis, depicting the percent change in metabolite levels (red line) and the line-broadening parameter sigma (black line) with their respective 95% confidence intervals (CI) indicated by the shaded regions. The onset and offset of the 10% and 100% STIM blocks are marked by dotted lines.

#### Sub-block model

When modeling the ON-OFF blocks during the STIM block, we found significant line narrowing for both the 10% and 100% contrasts (10%: z=-2.84, p=0.001; 100%: z=-3.24, p<0.001), but no difference between the contrasts (z=-1.18, p=0.12). We here report the results from the variable linewidth model, with the fixed linewidth model results in Supplementary Table 3. Visual stimulation significantly increased Glu levels for the 10% and 100% contrasts (10%: z=1.99, p=0.02; 100%: z=2.94, p=0.002), and across both contrasts (z=2.87, p=0.002). No effect of contrast was found on Glu levels (z=0.24, p=0.40). Additionally, increased tCr levels for the 100% contrast (z=2.26, p=0.01) and mean activation (z=2.25, p=0.01) were observed. These results are displayed in Figure 3 and Supplementary Figure 3. All statistical results can be found in Table 2.

**Figure 3.**
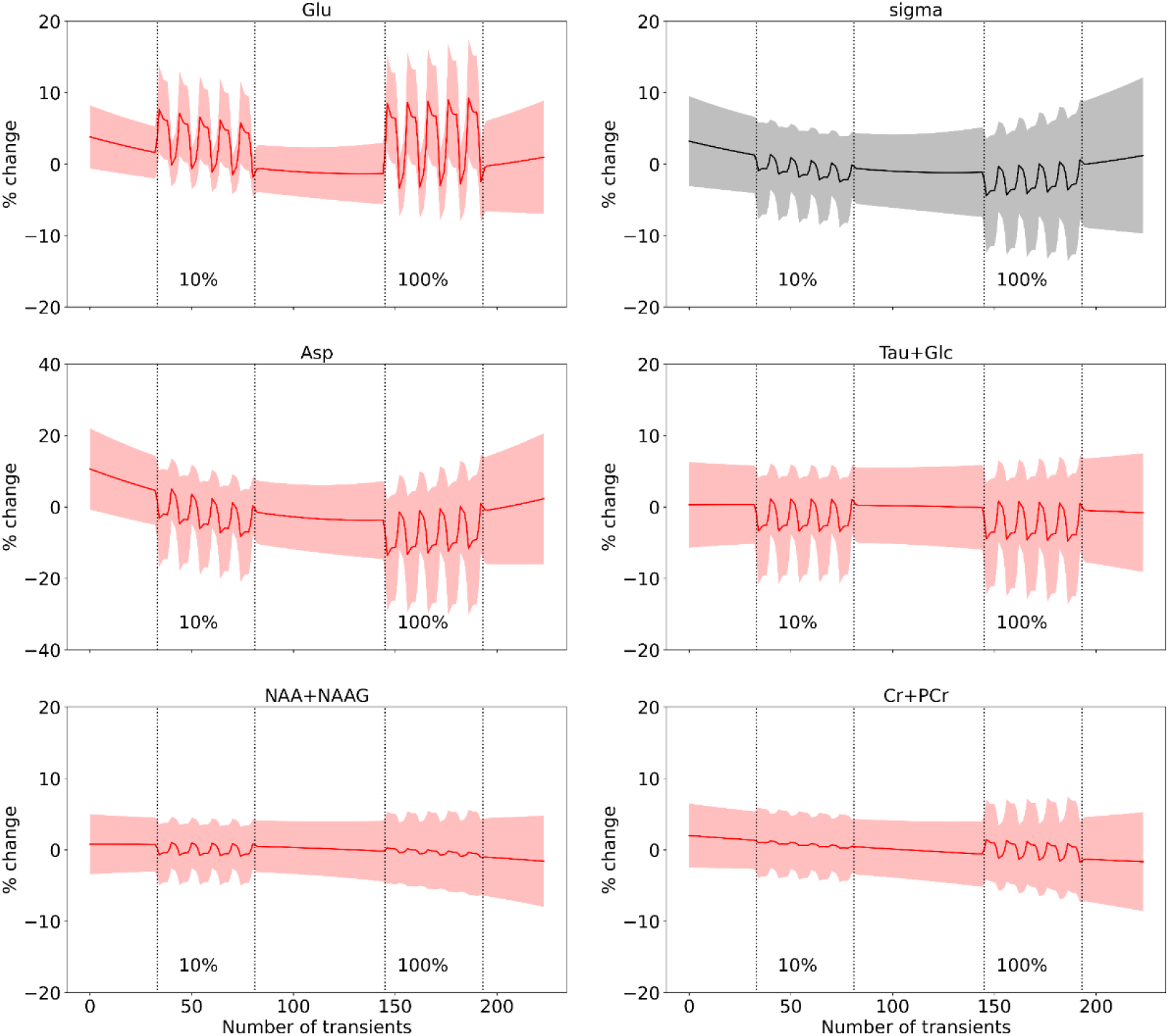
fMRS sub-block results. The figure displays significant results from the second-level fMRS analysis, depicting the percentage change in metabolite levels (red line) and the line-broadening parameter sigma (black line) with their respective 95% confidence intervals (CI) indicated by the shaded regions. The onset and offset of the 10% and 100% STIM blocks are marked by dotted lines.

### fMRI

Both the 10% and 100% contrast induced significant BOLD signal changes within the MRS voxel (10%: t=9.17; p<0.01; 100%: t=14.50; p<0.01) (Figure 4C), and we observed a higher increase in BOLD in the 100% contrast compared to the 10% contrast (100%>10%: t=8.70, p<0.01). This can also be observed in the whole brain analyses (Figure 4A).

**Figure 4.**
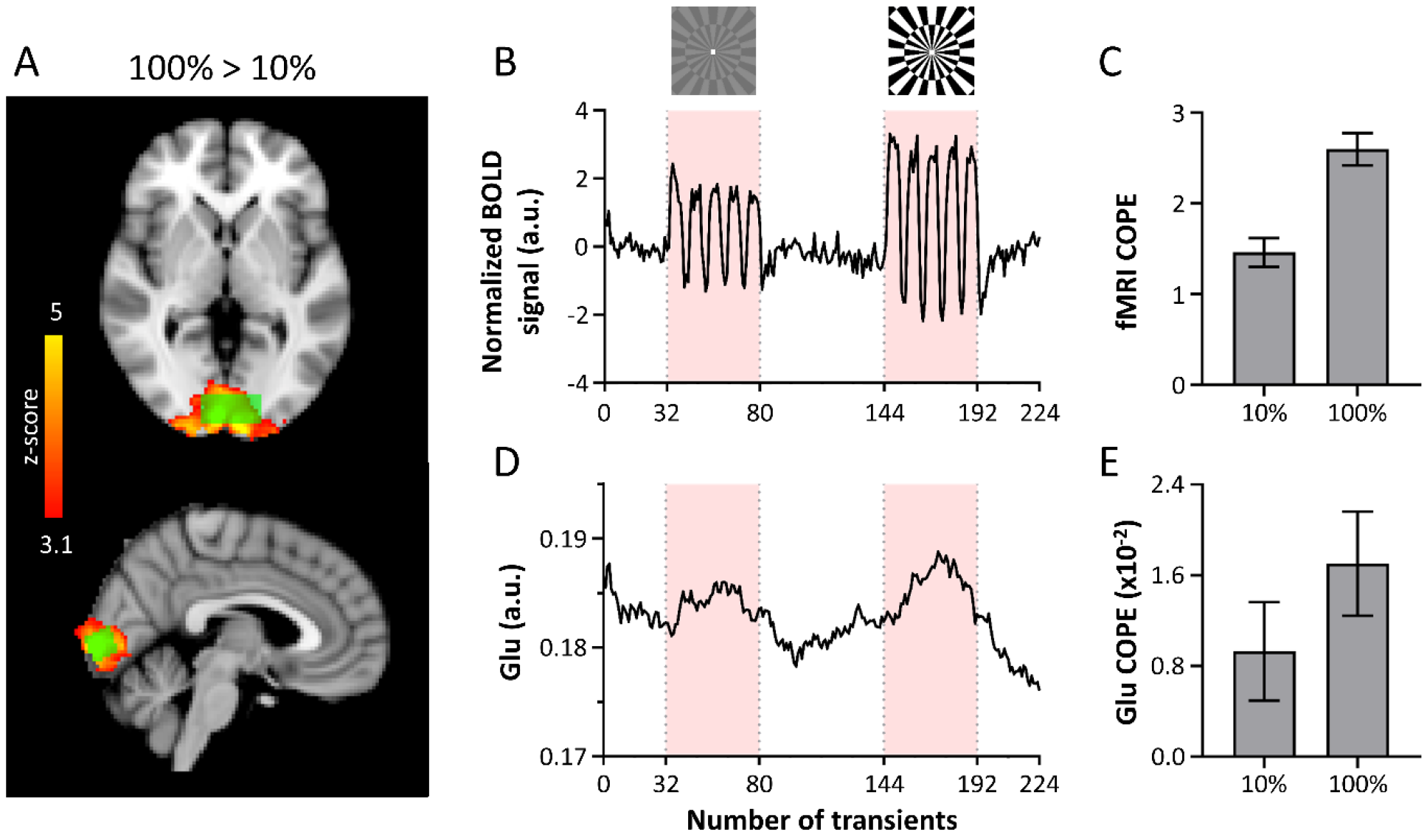
fMRI results. A) The z-map of the 100%>10% contrast (red/yellow) and a representative voxel (green) are overlaid on the MNI template. B) The mean BOLD signal of each individual’s MRS voxel was extracted and normalized to the mean of the first REST period. This graph shows the mean normalized time course across all subjects. C) Mean ± SEM BOLD signal COPE within the MRS voxel for the 10% and 100% contrast. D) The mean moving average (bin size = 32) of the initial (single transient) fits for Glu over time across subjects (extracted from FSL-MRS) E) Mean ± SEM Glu COPE within the MRS voxel for the 10% and 100% contrast. COPE = contrast parameter estimate

### Association between fMRS and fMRI response

The temporal signal envelope of the fMRS and fMRI data is visualized in Figure 4B and 4D. We plotted the contrast dependency of both the Glu and fMRI response in Figure 5 and show a non-linear relation with visual contrast. We did not find any significant correlations between the interindividual parameter estimates of the BOLD signal (fMRI) and Glu levels (fMRS), nor of the BOLD signal (fMRI) and the linewidth (fMRS). This outcome was consistent for both the full-block and sub-block analyses (Supplementary Figure 4; Supplementary Table 4).

**Figure 5.**
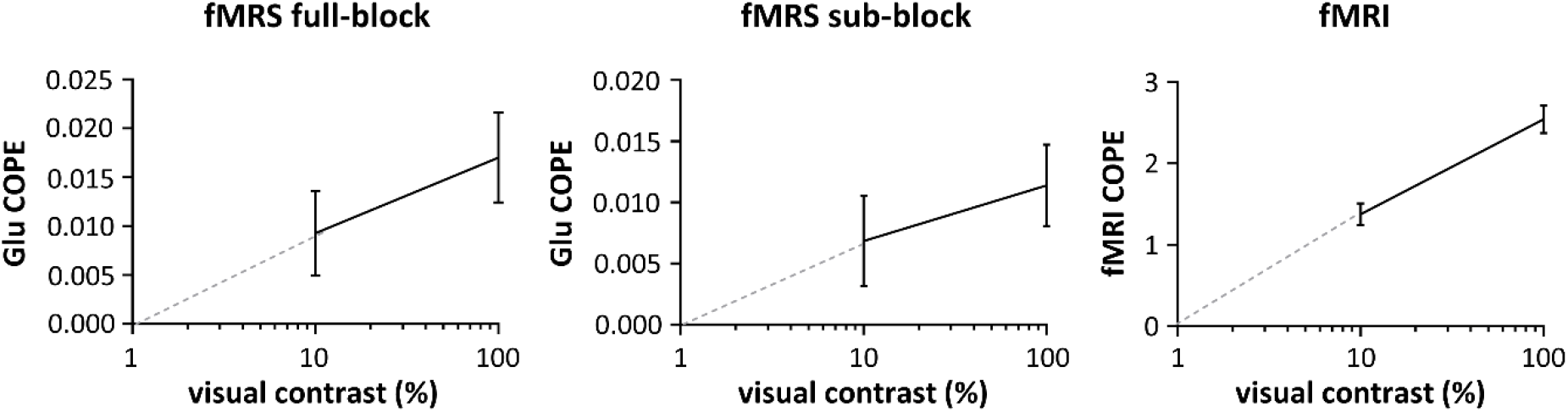
Non-linear fMRS and fMRI response to visual contrast. Graphs show the mean and standard error of the mean of first-level (individual) contrast parameter estimates (COPE) for A) Glu analyzed using full-block design B) Glu analyzed using sub-block design C) BOLD signal analyzed using sub-block design. The x-axis is plotted on a logarithmic scale. The gray dotted line shows the extrapolation to 0% contrast. The fMRS and fMRI response increases non-linearly with visual contrast.

## Discussion

### Metabolite response is not linearly related to visual contrast

We observed significantly higher BOLD activation for the 100% contrast compared to the 10% contrast (Figure 5C). This finding aligns with literature showing that the fMRI response, which is related to neural activity via a linear transform, exhibits a nonlinear response to visual contrast (Boynton et al., 1996; Logothetis et al., 2001). This is likely due to the nonlinearity of neural activity in V1, as suggested by Boynton et al. (1996). Interestingly, we observe a similar non-linear relationship between visual contrast and the neurometabolic response of glutamate, as illustrated in Figure 5A and B.

Our results partially align with one previous study investigating the neurometabolite response to visual contrast (Ip et al., 2019), in the sense that we also observe a monotonic relation between visual contrast and the fMRS response. However, while we find significant Glu increases at 10% visual contrast, they report no significant Glu increases for contrasts lower than 100% (i.e., 3%, 12.5%, and 50%). This could potentially be explained by differences in terms of stimuli timings and analysis methods. Nevertheless, we were also not able to discern significant differences between the 10% vs 100% contrast, suggesting that Glu changes in V1 in response to differences in visual contrast are subtle and challenging to detect, thus requiring a larger sample.

The neurometabolic demands of different visual stimuli have been investigated in previous studies. Bednarik et al. (2018) presented chromatic and achromatic stimuli that activate two separate clusters of neuronal populations, known as “blobs” and “interblobs,” that show comparable BOLD responses, but have different capacities for glucose oxidation. Despite their hypothesis, they did not observe any difference in neurometabolic responses elicited by the different stimuli. Another study investigated isoluminant chromatic stimuli at different flickering frequencies that are perceived and unperceived, but elicit comparable BOLD responses (diNuzzo et al., 2022). They observed markedly different neurometabolic responses, such that lactate and glutamate increased only when the flickering was perceived, but not when unperceived. Our study adds to the evidence that changes in neurometabolism are particularly sensitive to the type of information processing taking place in V1, rather than specifically targeted neuronal subpopulations. Therefore, fMRS can provide complementary information about cortical processing, in addition to BOLD fMRI.

Our study revealed that visual stimulation elicited changes in multiple other neurometabolites, including Asp and Glc+, which have been previously reported in the context of neural activation (*see* Koush et al. 2022). Consistent with prior research, we observed a decrease in these metabolites following visual stimulation, although the effect was more consistent across visual contrasts for the full block analysis. We found no significant differences in the change in Asp and Glc levels between the 10% and 100% contrasts. Reduced Glc levels are typically interpreted as the result of increased CMR_Glc_ in response to increased energy demands, while the opposite changes in Asp and Glu are thought to reflect an increase in the malate– aspartate shuttle (McKenna et al., 2006; Mangia et al., 2007). Furthermore, our study identified changes in neurometabolites that are less often reported, such as alterations tNAA and tCr. A previous study using functional diffusion-weighted MRS ascribed changes in tCr and tNAA diffusion properties to modifications in neuronal microstructure during neural activation, as well as to an increase in the energy-dependent cytoplasmic streaming associated with enhanced metabolism during visual stimulation (Branzoli et al., 2013, Sappey Marinier et al., 1992). Alternatively, given that changes in these metabolites tend to be observed mostly without correction for BOLD-induced linewidth changes (Mangia et al. 2006), these effects may be due to the incomplete accounting for T2* effects by the variable linewidth model.

### Comparison of full-block and sub-block analysis for fMRS

The characteristics of the BOLD fMRI response to visual stimulation have been extensively studied and are well documented (Jorge et al., 2018). Visual fMRI experiments commonly employ block durations of tens of seconds, eliciting a BOLD response following the hemodynamic response function. Indeed, our fMRI results showed a strong BOLD response that closely followed the experimental paradigm. However, in the case of fMRS, the existence of an analogous ‘neurometabolic response function’ remains uncertain. To further investigate this temporal response, we examined whether a sustained response was present throughout the entire stimulus block (full-block analysis) or if there was a more rapid response akin to fMRI (sub-block analysis).

Our results indicate that the full-block model may capture the data slightly better as compared to the sub-block model, as evidenced by stronger effects for the neurometabolites (z-scores) and better fits of the model (lower AICs). This suggests that the paradigm may have elicited mostly sustained changes in neurometabolite levels. As such, modeling the OFF blocks in between the ON blocks may have increased the residuals of predicted-to-measured neurometabolite changes. This is supported by results from an animal fMRS study employing a comparable design (Ligneul et al. 2021). They also used stimulus blocks with an ON-OFF design, but with a longer OFF period compared to our study, and while they found evidence for sustained increase in Glu during the stimulation blocks, they did not detect faster neurometabolic responses following the ON-OFF design.

Nevertheless, it is plausible that different neurometabolites may exhibit dissimilar temporal response profiles. Certain neurometabolites may demonstrate fluctuations at faster temporal scales, while others may manifest changes only following sustained stimulation. This would be dependent upon the respective neurometabolite’s function in neuronal energetics and neurotransmission. Insights from mathematical modeling have provided evidence for both rapid as well as prolonged changes in neurometabolites. For example, sustained neurometabolite changes have been attributed to increased activity in metabolic pathways, i.e. via neurometabolite synthesis (Mangia et al., 2009) whereas recent work has suggested that compartmental shifts of metabolites could explain more rapid changes in neurometabolites during fMRS experiments (Lea-Carnall et al. 2023). Future studies should focus on experiments teasing apart the contribution of different cellular compartments to the overall metabolite signal, to characterize the ‘neurometabolite response function’. In addition, future advancements in dynamic fitting models could conceivably integrate varied models to account for the idiosyncratic temporal profiles of different neurometabolites and hemodynamic responses.

### Analysis considerations for fMRS data: dynamic fitting and linewidth effects

The data in this study were analyzed using a novel dynamic fitting algorithm implemented in FSL-MRS (Clarke et al., 2023). While spectral-temporal or GLM analysis has been employed previously (Tal 2022, Ligneul et al., 2021), most previous studies have utilized block averaging to demonstrate neurometabolic changes in fMRS experiments. Compared to block averaging, GLMs can potentially increase sensitivity and specificity in detecting neurometabolite alterations associated with experimental conditions, even in the presence of noise and variability, because they allow for more comprehensive modeling of the fMRS temporal signal (Tal et al., 2023, Clarke et al., 2023). It takes into account the temporal dynamics of the fMRS signal, rather than collapsing the temporal dimension of the data. Nonetheless, the accuracy of the model and design matrix that we chose for the GLM is crucial for the accuracy and validity of the results obtained from these analyses. Importantly, as mentioned above, we currently do not know the properties of the neurometabolite response function(s). Future studies are needed to experimentally validate the precise specification of the statistical model, which includes the selection and definition of design variables, the estimation of model parameters, and the assumptions underlying the model.

In light of previous investigations illustrating that local magnetic field alterations provoked by neural activity can induce linewidth alterations (Zhu & Chen, 2001; Mangia et al., 2006), we assessed the effect of incorporating a variable (Gaussian) linewidth parameter into our dynamic model. Our analysis revealed that permitting the linewidth to fluctuate significantly contributed to the model for both the full-block and sub-block approaches. Comparatively, the outcomes stemming from the fixed linewidth model (Supplements) demonstrated more significant results for the neurometabolites than those associated with a variable linewidth model. For example, significant differences in the tNAA and tCr resonances were found to be partially accounted for by adding the dynamic linewidth parameter. This corroborates block-averaging studies (e.g. Mangia et al., 2007), which underscore the necessity of correcting for linewidth to prevent false positives.

One limitation of this work is that we consistently presented the 10% contrast prior to the 100% contrast. This was chosen in order to prevent potential spillover effects from the 100% contrast. Another limitation is that, based on the quality of the spectra, we could not interpret the results of Ala, GABA, Gln and Lac statistically, even though these metabolites are of interest due to their role in neurometabolism. In contrast to our expectations and previous findings (e.g. Bednarik et al. 2015), we did not find a negative association between the BOLD effects as estimated by fMRI and linewidth changes in fMRS. This may be due to the large interindividual variation in the sigma estimates compared to the BOLD estimates.

## Conclusion

In conclusion, our study provides evidence of a nonlinear relationship between visual contrast and both the BOLD response and the glutamate response in V1. In line with previous literature, we also identified concomitant changes in several other neurometabolites, including Asp, Glc, Tau, tNAA, and tCr. Moreover, our dynamic fitting approach allowed us to compare both sustained and more rapid neurometabolite responses. Although the data are suggestive of an overall more sustained response, future studies should explore the temporal response profiles of different neurometabolites and further refine the statistical models used for fMRS analysis. Notwithstanding its limitations, our study provides valuable insights into the complex relationship between visual contrast, neural activity, and neurometabolism, highlighting fMRS as a complementary technique to BOLD fMRI.

## Acknowledgements

The authors thank Dr Vincent Boer for his help with the data acquisition software.

