## Supplementary Materials for "A 7T interleaved fMRS and fMRI study on visual contrast dependency in the human brain"

### Supplementary Material

**Supplementary Table 1. MRSinMRS checklist.** Description of MRS data acquisition, analysis, and quality assessment.

| **MRSinMRS checklist** | |
|---|---|
| **1. Hardware** | |
| **a. Field Strength** | 7T |
| **b. Manufacturer** | Philips |
| **c. Model** | Achieva |
| **d. RF coil** | Quadrature transmit/32-channel receive head coil (Nova Medical) |
| **e. Additional hardware** | Dielectric pad or metasurface (Webb et al. 2022) |
| **2. Acquisition** | |
| **a. Pulse sequence** | Interleaved sLASER and 3D-EPI |
| **b. Volume of interest** | V1 |
| **c. Nominal VOI size** | 14 x 31 x 14 mm |
| **d. Repetition time (TR), echo time (TE)** | TR/TE=3600/36 ms |
| **e. Total number of acquisitions/averages** | 224 |
| **i. Number of averaged spectra per time point** | n.a. |
| **ii. Averaging method** | None, dynamic spectral-temporal fitting |
| **f. Additional sequence parameters** | 3000 Hz; 1024 data points |
| **g. Water suppression method** | VAPOR |
| **h. Shimming method** | HOS-DLT (Boer et al. 2020) |
| **i. Triggering or motion correction method** | None |
| **3. Data analysis methods and outputs** | |
| **a. Analysis software** | FSL-MRS v2.1.6 |
| **b. Processing steps deviating from quoted product** | Processing using in-house Matlab scripts (coil combine, ecc, spectral registration) |
| **c. Output measure** | Dynamic fitting on unscaled spectra |
| **d. Quantification references and assumptions** | Default basis set |
| **4. Data quality** | |
| **a. Reported variables** | SNR, FWHM |
| **b. Data exclusion criteria[1]** | Metabolites not evaluated when >50% subjects >20% CRLB |
| **c Quality measures of post processing model fitting** | Glu CRLB, NAA FWHM |
| **d. Sample spectrum** | Figure 1 |

**Supplementary Figure 1. The specified models in FSL-MRS**

Fixed model

```
# Parameter - functional relationships
Parameters = {
'conc'     : {'dynamic':'model_glm','params':[f'beta{i}' for i in range(5)]},
'gamma'    : 'fixed',
'sigma'    : 'fixed',
'eps'      : 'fixed',
'baseline' : 'fixed',
'Phi_0'    : 'fixed',
'Phi_1'    : 'fixed'
}

# Bounds on free fitted parameters
Bounds = {
'gamma' : (0,None),
'beta4' : (0,None),
}

# Dynamic models
from numpy import dot
def model_glm(p,t):
    return dot(t,p)

# Dynamic model gradients
def model_glm_grad(p,t):
    return t.T
```

Variable linewidth model full-block

```
# Parameter - functional relationships
Parameters = {
'conc'     : {'dynamic':'model_glm_conc','params':['stim0', 'stim1', 'drift_1', 'drift_2', 'constant']},
'gamma'    : 'fixed',
'sigma'    : {'dynamic':'model_glm_sigma','params':['stim0', 'stim1', 'drift_1', 'drift_2', 'constant']},
'eps'      : 'fixed',
'baseline' : 'fixed',
'Phi_0'    : 'fixed',
'Phi_1'    : 'fixed'
}

# Bounds on free fitted parameters
Bounds = {
'gamma' : (0,None),
'constant': (0, None)}


# Dynamic models
#from numpy import dot
#def model_glm(p,t):
#    return dot(t,p)

# Dynamic model gradients
#def model_glm_grad(p,t):
#    return t.T

# Dynamic models
from numpy import dot

def model_glm_conc(p, t):
    return dot(t[:, [0, 1, 4, 5, 6]], p)

# Dynamic model gradients
def model_glm_conc_grad(p, t):
    return t[:, [0, 1, 4, 5, 6]].T

# Dynamic models
def model_glm_sigma(p, t):
    return dot(t[:, [2, 3, 4, 5, 6]], p)

# Dynamic model gradients
def model_glm_sigma_grad(p, t):
    return t[:, [2, 3, 4, 5, 6]].T
```

### Variable linewidth model sub-block

```
# Parameter - functional relationships
Parameters = {
'conc'     : {'dynamic':'model_glm','params':[f'beta{i}' for i in range(5)]},
'gamma'    : 'fixed',
'sigma'    : {'dynamic':'model_glm','params':[f'beta{i}' for i in range(5)]},
'eps'      : 'fixed',
'baseline' : 'fixed',
'Phi_0'    : 'fixed',
'Phi_1'    : 'fixed'
}

# Bounds on free fitted parameters
Bounds = {
'gamma' : (0,None),
'beta4' : (0,None),
}

# Dynamic models
from numpy import dot
def model_glm(p,t):
    return dot(t,p)

# Dynamic model gradients
def model_glm_grad(p,t):
    return t.T
```

### Supplementary Table 2. Statistical results for the fixed linewidth full-block analysis.

| | 10% | | 100% | | mean activation | | 100 vs 10% | |
|---|---|---|---|---|---|---|---|---|
| | *z* | *p* | *z* | *p* | *z* | *p* | *z* | *p* |
| **Metabolite level** | | | | | | | | |
| *Asc* | 1.01 | 0.16 | 0.32 | 0.38 | 0.62 | 0.27 | -0.53 | 0.30 |
| *Asp* | -1.69 | 0.046 | **-2.04** | **0.02** | **-2.23** | **0.01** | -0.42 | 0.34 |
| *GSH* | 0.08 | 0.47 | 1.48 | 0.07 | 1.06 | 0.15 | 1.08 | 0.14 |
| *Glu* | **2.39** | **0.01** | **3.56** | **<0.001** | **3.45** | **<0.001** | 1.35 | 0.09 |
| *Ins* | -0.37 | 0.36 | **2.14** | **0.02** | 0.14 | 0.86 | 1.62 | 0.053 |
| *PE* | 0.24 | 0.40 | -0.84 | 0.20 | 0.92 | 0.08 | 1.19 | 0.12 |
| *Scyllo* | 0.36 | 0.36 | 0.11 | 0.46 | 0.38 | 0.62 | -0.19 | 0.43 |
| *Glc+* | -1.80 | 0.04 | **-2.43** | **0.008** | **-2.85** | **0.002** | -0.39 | 0.35 |
| *tCh* | 1.55 | 0.06 | -0.39 | 0.35 | 0.80 | 0.21 | 1.42 | 0.08 |
| *tCr* | 0.15 | 0.44 | **2.50** | **0.01** | **2.30** | **0.01** | 1.66 | 0.049 |
| *tNAA* | **-2.09** | **0.02** | 0.97 | 0.17 | -0.89 | 0.19 | 1.70 | 0.04 |

*Positive and negative z-values indicate a neurometabolite increase and decrease, respectively, in case of the 10%, 100% and mean activation. For the contrast between 10% vs 100% a positive z-value indicates 100% > 10% , whereas a negative z-value indicates 10% > 100%. The significant effects (p<0.05) are displayed in bold.*

### Supplementary Table 3. Statistical results for the fixed linewidth sub-block analysis.

| | 10% | | 100% | | mean activation | | 100 vs 10% | |
|---|---|---|---|---|---|---|---|---|
| | *z* | *p* | *z* | *p* | *z* | *p* | *z* | *p* |
| **Metabolite level** | | | | | | | | |
| *Asc* | 1.13 | 0.13 | 0.23 | 0.41 | 0.76 | 0.22 | -0.62 | 0.27 |
| *Asp* | -1.24 | 0.11 | -1.40 | 0.08 | -1.72 | 0.04 | -0.08 | 0.47 |
| *GSH* | 0.73 | 0.23 | 1.54 | 0.06 | **-3.04** | **0.001** | 0.62 | 0.27 |
| *Glu* | **2.25** | **0.01** | **3.64** | **<0.001** | **3.28** | **0.001** | 0.62 | 0.27 |
| *Ins* | 0.35 | 0.36 | **2.31** | **0.01** | 1.77 | 0.04 | 1.13 | 0.13 |
| *PE* | -1.48 | 0.07 | 0.43 | 0.34 | -1.22 | 0.11 | 1.14 | 0.13 |
| *Scyllo* | -0.57 | 0.29 | 0.23 | 0.41 | -0.24 | 0.41 | 0.57 | 0.28 |
| *Glc+* | -1.70 | 0.04 | -1.56 | 0.06 | **-2.39** | **0.008** | 0.31 | 0.38 |
| *tCh* | **2.70** | **0.004** | 1.08 | 0.14 | **2.56** | **0.01** | -1.34 | 0.91 |
| *tCr* | 1.18 | 0.12 | **3.18** | **0.001** | **3.30** | **<0.001** | 1.59 | 0.06 |
| *tNAA* | -0.55 | 0.29 | 1.34 | 0.09 | 0.96 | 0.17 | 1.24 | 0.11 |

*Positive and negative z-values indicate a neurometabolite increase and decrease, respectively, in case of the 10%, 100% and mean activation. For the contrast between 10% vs 100% a positive z-value indicates 100% > 10% , whereas a negative z-value indicates 10% > 100%. The significant effects (p<0.05) are displayed in bold.*

**Supplementary Figure 2B. Individual and mean traces of full-block analysis**

Individual level traces extracted from first-level analysis are shown in red with the mean trace across subjects extracted from the second-level analysis in black. For visualization purposes, y-axes are cut-off and do not include the full extent of the plotted individual trace.

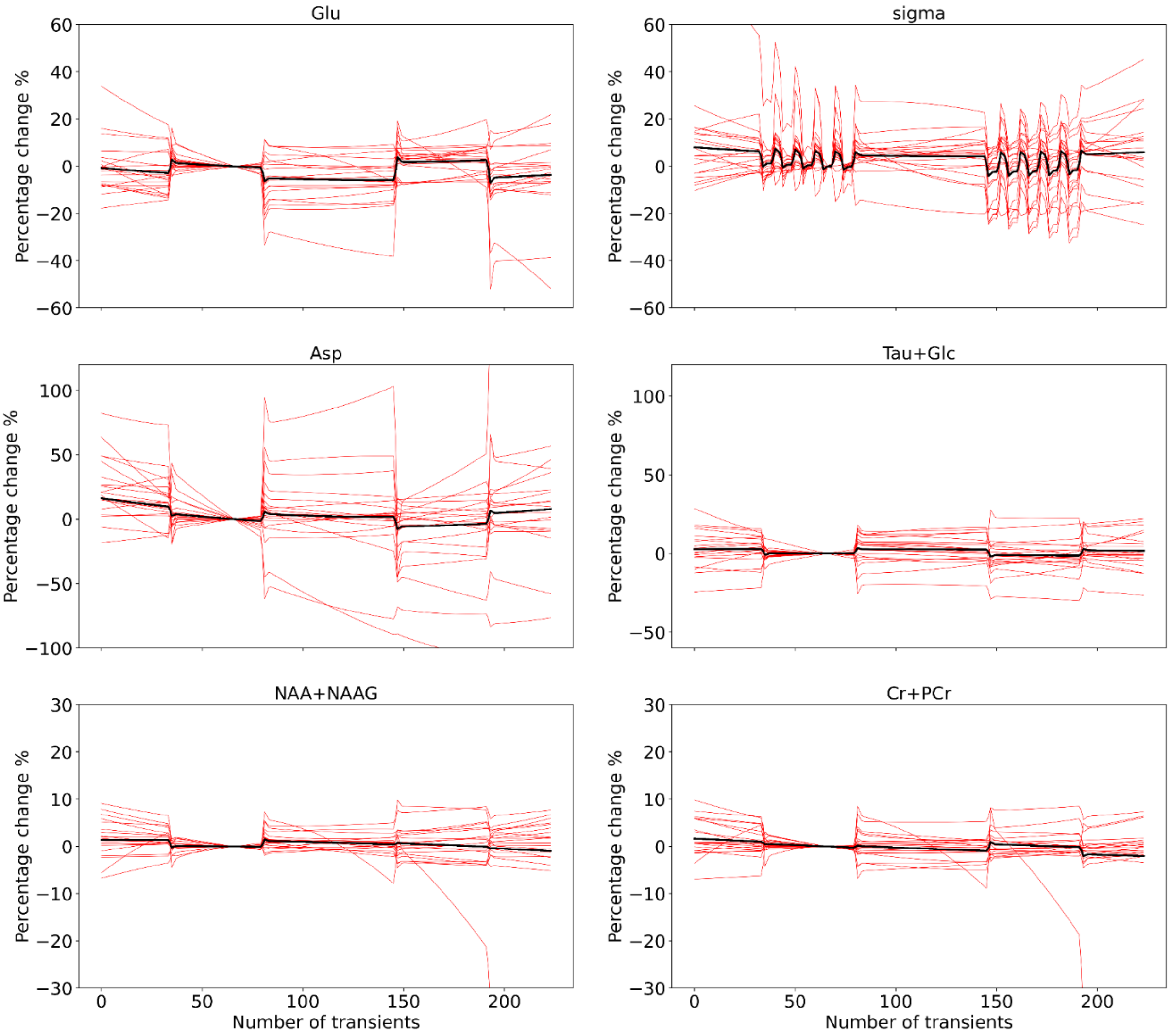

**Supplementary Figure 2B. Individual and mean traces of full-block analysis**

Same as 2A, but with specifically only showing the task effect, i.e. with the drift and constant terms removed.

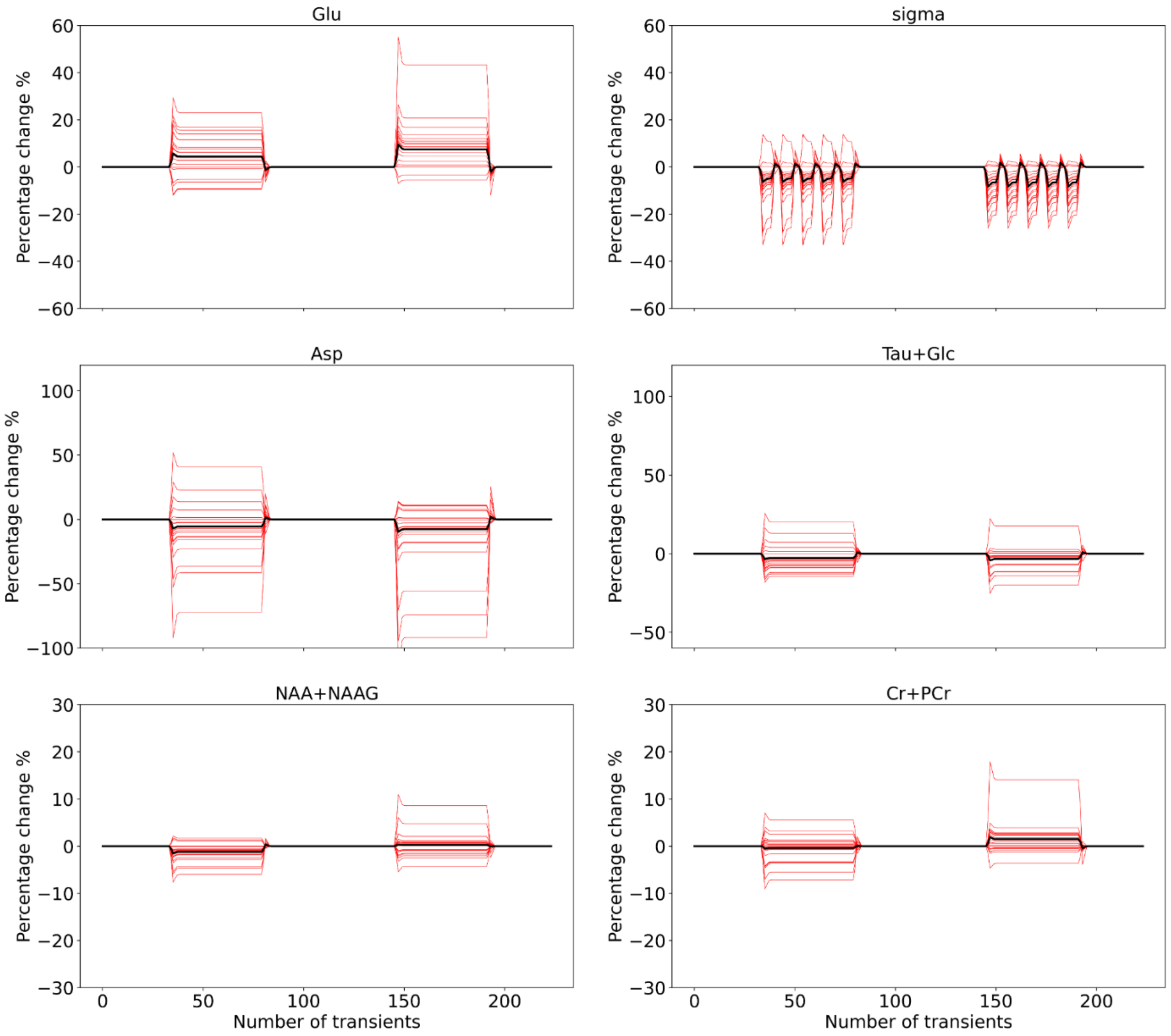

**Supplementary Figure 3A. Individual and mean traces of sub-block analysis**

Individual level traces extracted from first-level analysis are shown in red with the mean trace across subjects extracted from the second-level analysis in black. For visualization purposes, y-axes are cut-off and do not include the full extent of the plotted individual trace.

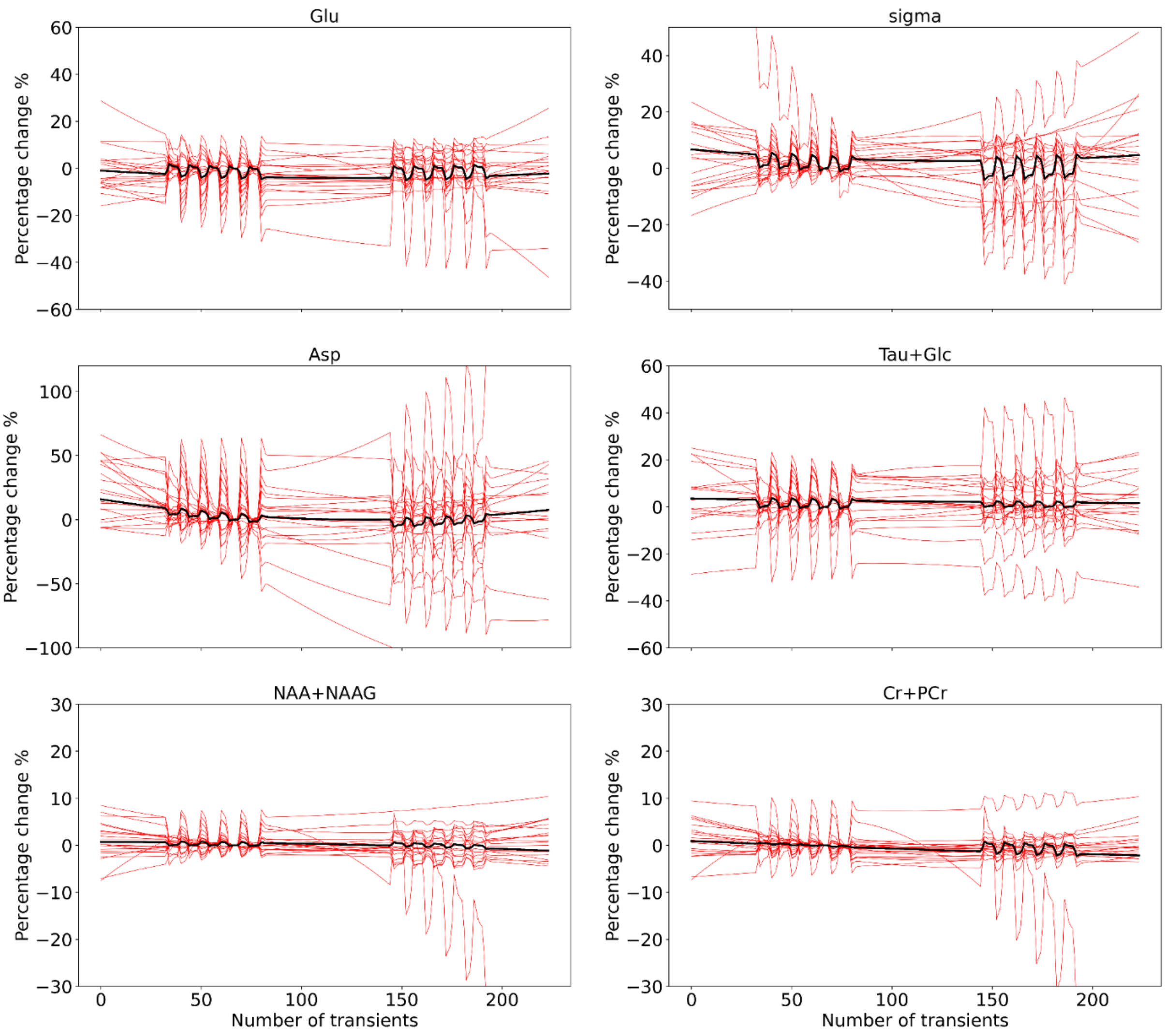

**Supplementary Figure 3B. Individual and mean traces of sub-block analysis**

Same as 3A, but with specifically only showing the task effect, i.e. with the drift and constant terms removed.

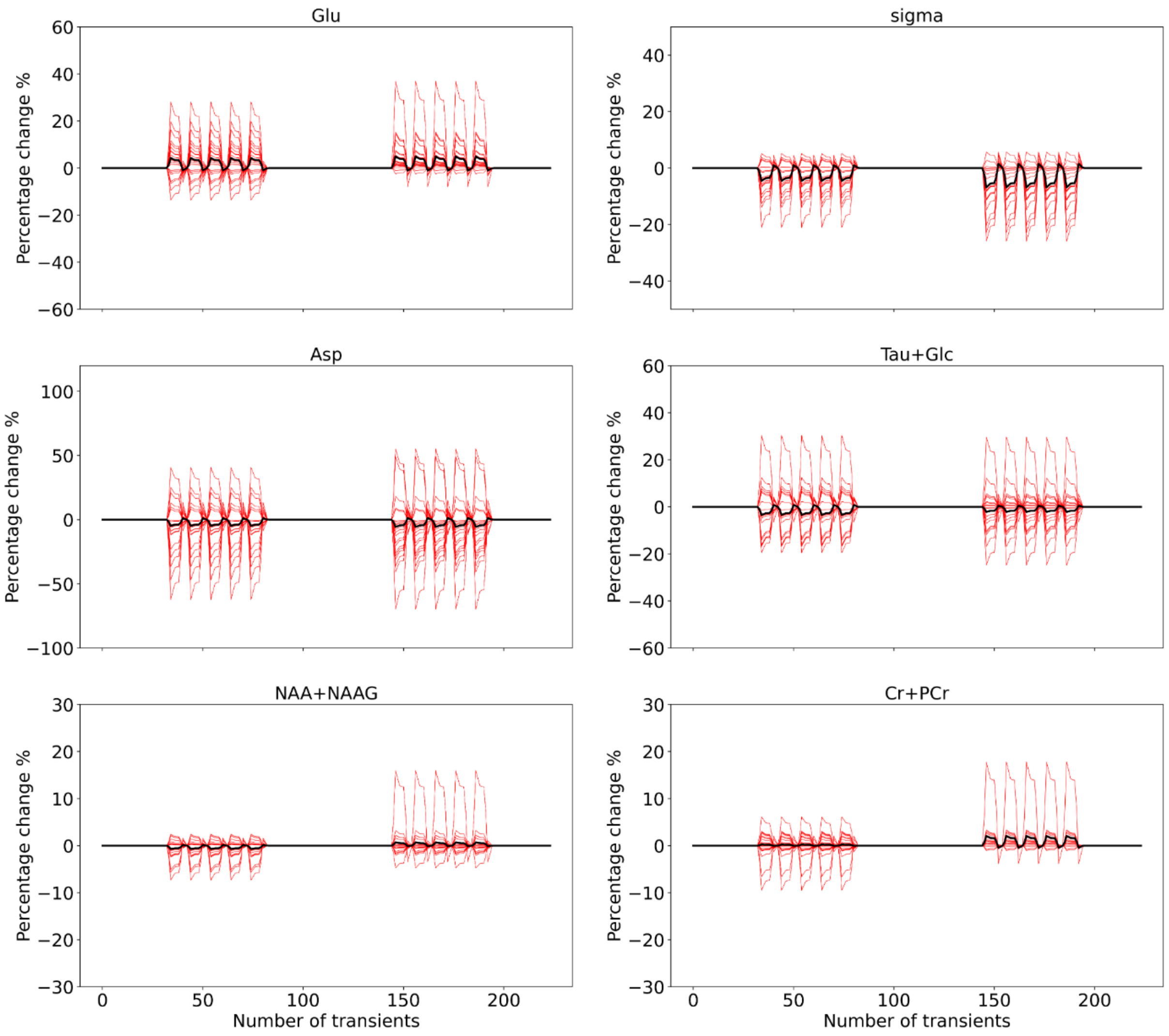


**Supplementary Table 4. Correlations between fMRI and fMRS**

| | 10% | | 100% | | 100 vs 10% | |
|---|---|---|---|---|---|---|
| | *r* | *p* | *r* | *p* | *r* | *p* |
| **Full-block** | | | | | | |
| *fMRI BOLD vs fMRS Glu* | 0.49 | 0.027 | -0.006 | 0.98 | 0.23 | 0.33 |
| *fMRI BOLD vs fMRS sigma* | -0.07 | 0.74 | -0.25 | 0.29 | 0.007 | 0.97 |
| **Sub-block** | | | | | | |
| *fMRI BOLD vs fMRS Glu* | 0.49 | 0.03 | 0.07 | 0.78 | 0.45 | 0.05 |
| *fMRI BOLD vs fMRS sigma* | -0.25 | 0.28 | 0.47 | 0.03 | 0.31 | 0.19 |

**Supplementary Figure 4. Correlations between fMRI and fMRS**

Scatterplots of the fMRI BOLD signal (x-axis) and the glutamate (top rows) or the sigma, i.e. the linewidth changes (bottom rows) for the 10%, 100% and 100>10% conditions. A linear regression line through the individual data points is shown with the 95% confidence intervals in dotted lines.

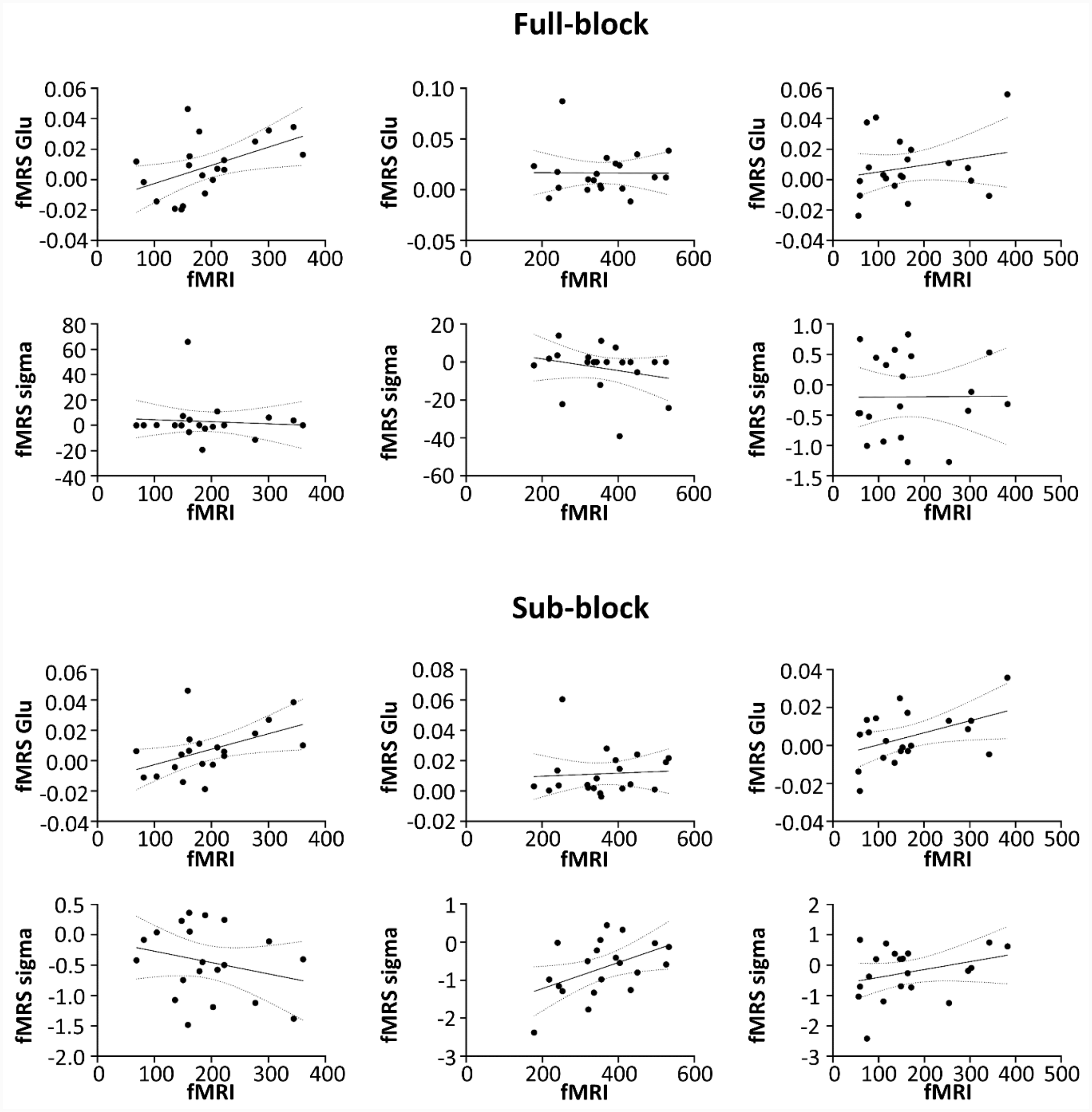
